# A Modular Platform for Effector Discovery in Induced-Proximity Lysine Acetylation

**DOI:** 10.64898/2026.03.11.711209

**Authors:** Brianna Hill-Payne, Mohd. Younis Bhat, George M. Burslem

## Abstract

The regulation of post-translational modifications (PTMs) is central to cellular biology and disease. Induced-proximity strategies enable manipulation of PTMs by recruiting modifying enzymes to proteins of interest, but identifying effective effector enzymes typically requires extensive heterobifunctional molecule synthesis before biological validation. Here we report a modular platform that enables rapid evaluation of PTM editing enzymes against defined protein substrates in living cells using compound-dependent or nanobody-mediated induced proximity. Using lysine acetylation as a model system, we demonstrate programmable acetylation of GFP, histone H3, and p53 through recruitment of diverse acetyltransferases. Effector identity dictates site-specific acetylation patterns, enabling selective PTM deposition across substrates and cellular compartments. This platform enables rapid identification of productive effector–substrate relationships prior to heterobifunctional molecule development, accelerating the design of induced-proximity chemical probes for targeted PTM editing.

## Introduction

Protein homeostasis is essential for cell survival, yet it is a complex network involving a multitude of key players. Post-translational modifications (PTMs) are central regulators that dynamically expand proteomic function without requiring de novo protein synthesis ^1–4^. Biochemical tools that manipulate PTMs continue to expand, ranging from broad high-throughput methods for enzyme identification to precise amino acid changes^5–13^. Whether for therapeutic design or mechanism determination, the ability to control a protein’s PTM status continues to be widely sought after. Recent advances in chemically induced proximity have enabled protein-specific manipulation of PTMs in living cells ^7, 8, 13–18^.

Induced proximity has recently developed into an expansive area of research that enables the investigation of specific proteins of interest while minimizing perturbations to the broader proteome ^14, 16, 19^. The use of heterobifunctional molecules, where two protein binding elements are joined by a linker, has proven effective in inducing the colocalization of two proteins ^7^. The most notable heterobifunctional molecules are proteolytic targeting chimeras (PROTACs), where a heterobifunctional molecule induces the localization of an E3 ligase to a protein of interest, triggering ubiquitination and subsequent proteasomal degradation ^7, 16, 20, 21^. The use of heterobifunctional molecules in PTM characterization has since expanded to other modifications with AceTAGs, DUBTACs, PHICs, and many others that involve localizing a PTM writer enzyme to a protein of interest^8, 9, 13, 22–26^.

Despite the rapid expansion of induced-proximity technologies, identifying effective effector enzymes for a given target protein remains largely empirical. In current workflows, heterobifunctional molecules are often designed before it is known whether the recruited enzyme can efficiently modify the intended substrate, leading to extensive synthetic effort prior to biological validation. High-throughput methods, including our own SPOTLITES methods, allow the identification of optimal enzymatic hits against a target protein by screening a pool of PTM writers ^5, 6, 27^. Such methods have created opportunities to identify ideal protein-enzyme interactions prior to engaging in extensive molecule design. However, these methods do not directly enable rapid validation of effector–substrate compatibility in living cells. As a result, a critical gap exists between large-scale effector discovery and the development of proximity-inducing chemical probes. A modular platform capable of rapidly evaluating PTM editors in defined induced-proximity configurations would therefore accelerate both mechanistic studies of PTM biology and the development of next-generation proximity-based chemical tools.

To address this challenge, we envisioned a modular system for the rapid testing of biological hypotheses and/or the small-scale exploration of which effectors may be the most useful within a family of PTM editors, including whether different effectors could result in different patterns of PTM deposition. By creating fusion proteins that harness the dimerization of HaloTag7 and FKBP12^F36V^ via a generic heterobifunctional molecule, HaloFK7, or via the application of a protein-specific nanobody, we created a modular system that can conceptually be used to effectively and efficiently induce localization of any PTM editing enzyme to any protein of interest ^5, 6, 28, 29^.

To validate our platform’s capabilities, we focused on its ability to induce targeted acetylation on a protein of interest. Acetylation, an essential PTM, plays a prominent role in protein homeostasis with direct and indirect implications across the proteome ^1, 2, 30, 31^. The reversible mechanism, catalyzed by lysine acetyltransferases (KATs) and deacetylases (KDACs), is paramount in both nuclear and cytosolic protein interactions, like histone acetylation which leads to the opening of chromatin or FSP1 acetylation which regulates ferroptosis, making it an ideal candidate to assess our platform’s capabilities ^32, 33^. Here, we demonstrate the modularity of our platform by rapidly creating a series of three-component mammalian-expression vectors that can induce targeted acetylation in both compound-dependent and -independent modes (Fig. 1A). We demonstrate that our modular strategy can be used to rapidly develop strategies to alter the acetylation status of a multitude of substrates (GFP, p53 and H3), via recruitment of various PTM editors from the acetyltransferase super family (p300, PAT, Tip60, GCN5), we show that site-selective acetylation patterns are dictated by the recruited enzyme rather than by the induced proximity. Collectively, this work establishes a programmable and modular PTM editing framework that bridges effector selection and heterobifunctional molecule development, enabling rapid interrogation of effector specificity, substrate compatibility, and xeno-enzyme deployment in mammalian cells. By decoupling effector discovery from chemical probe development, this approach provides a general framework for identifying functional PTM editing enzymes, exemplified with acetylation, prior to the design of heterobifunctional molecules or other proximity-inducing modalities.

**Figure 1:**
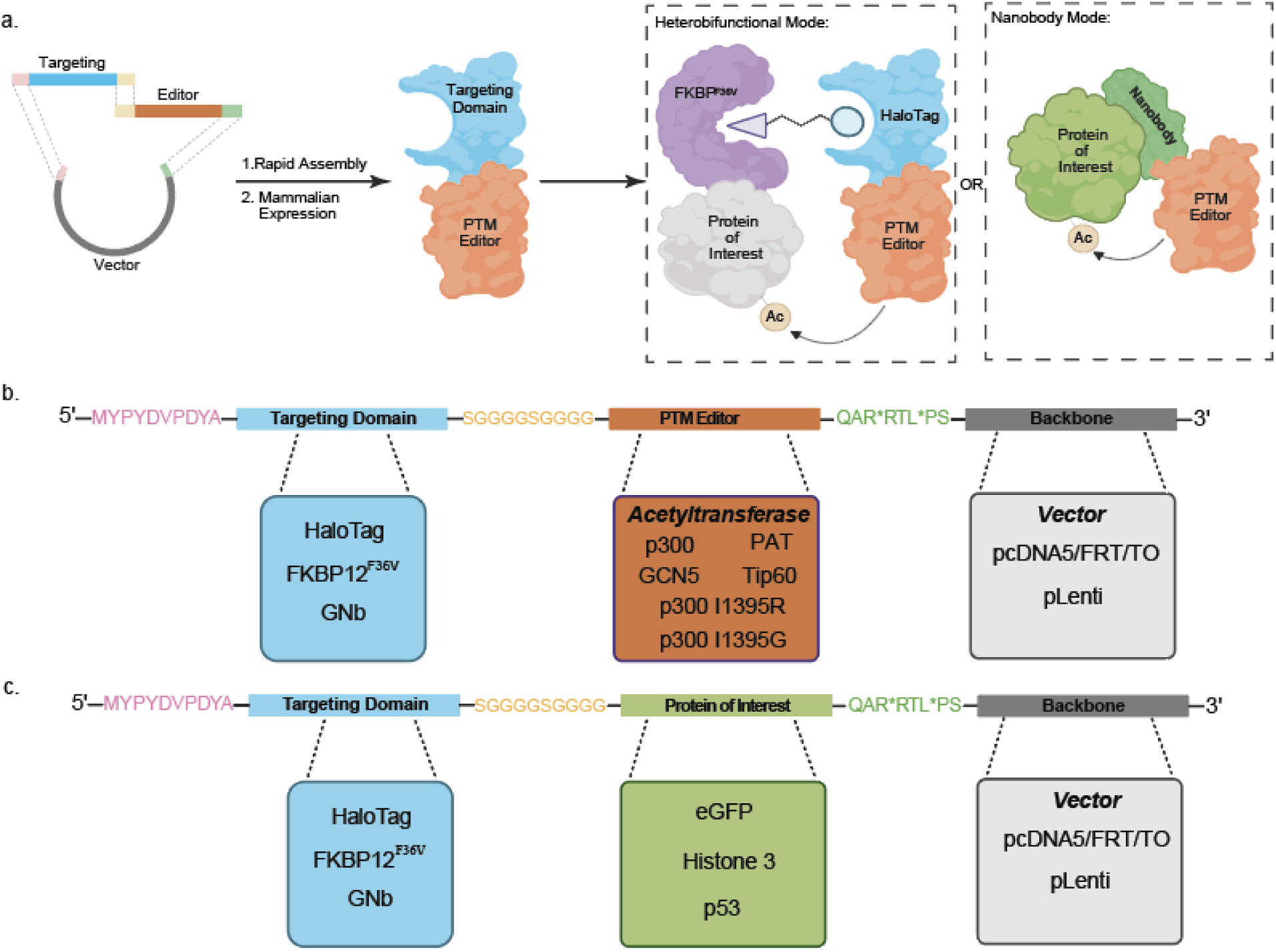
Induced Proximity Approach and Modular Cloning Strategy. A) Schematic representation of our modular approach for both heterobifunctional and nanobody modes of induced proximity. B) Gibson overhangs and components for rapid modular generation of targeting-effector fusion vectors. C) Gibson overhangs and components for rapid modular generation of protein of interest fusion vectors.

## Results & Discussion

### Generation of Modular PTM Editing Platform

Having set out to develop a platform to induce the targeted acetylation of a protein of interest in both heterobifunctional compound-dependent and -independent modes, we started by designing two fusion proteins with complementary ligandable domains, HaloTag7 and FKBP12^F36V,^ fused via a serine-glycine linker to either the protein of interest or the acetyltransferase (effector) ^22, 28, 34, 35^. HaloFK7, our previously reported heterobifunctional molecule induces the dimerization of HaloTag7 and FKBP12^F36V^ through covalent and non-covalent interactions, respectively, and thus can be used induce proximity between our fusion proteins in a heterobifunctional compound-dependent manner ^6^ (Fig.1A). Alternatively, one fusion protein where a protein-specific nanobody is fused to an acetyltransferase can be utilized for targeted acetylation in the absence of a heterobifunctional molecule (Fig. 1A). This platform is realized through a modular cloning strategy that enables the rapid creation of these fusion proteins. Utilizing Gibson assembly, a versatile and user-friendly technique (see detailed protocol in SI), we developed a three-part cloning strategy (targeting domain, effector domain, and backbone) to rapidly generate vectors containing our fusion proteins ^36^. The vectors can be generated for a variety of applications, such as bacterial expression, viral transduction, or mammalian overexpression, further diversifying user accessibility to elucidate biological mechanisms. The modularity of the platform is such that each component shares complementary overhangs required for a successful Gibson assembly and essential for retaining the interchangeability in constructs. After multiple rounds of optimization, we have verified that the modular overhangs utilized are robust and can successfully create a library of fusion proteins with varied targeting/effector domains, proteins of interest, and backbones (Fig 1B, 1C, SI Table 1). Whole plasmid sequencing was used to verify the successful assembly of each fusion protein vector created. Together, this modular cloning strategy provided a uniform method for constructing an ever-expanding collection of mammalian-expressing and lentiviral vectors, which is necessary for the realization of the platform.

### Proof of Concept - Targeted acetylation of eGFP by p300

To provide proof of concept for our strategy, we first assessed our ability to recapitulate the effect of recruiting the promiscuous acetyltransferase p300 to eGFP ^37–39^. Utilizing the cloning strategy described above, we generated a HA-HaloTag7-p300^1048–1660^ mammalian expression vector and transfected it into HEK293T cells stably transduced with a FKBP12^F36V^-GFP fusion construct. After confirming expression of HA-HaloTag-p300^1048–1660^ at the anticipated molecular weight (Fig. S1), we assessed the ability of HaloFK7 to induce a ternary complex between p300 and GFP (Fig. 2A). Notably, we observed the presence of HaloTag7-p300^1048–1660^ following GFP pulldown only when HaloFK7 is present, indicating that ternary complex formation is dependent on the heterobifunctional compound (Fig. 2B). Concurrent with ternary complex formation, we observed increased acetylation of eGFP in a compound-dependent fashion (Fig. 2C), and this could be further enhanced by co-treatment with a lysine deacetylase inhibitor (SAHA, 5 µM) (Fig 2D). Additionally, we observed clear dose-dependent ternary complex formation and the hook effect, another indicator of heterobifunctional compound-induced ternary complex formation (Fig. 2E). These results suggest that the induced proximity of HaloTag7-p300^1048–1660^ to FKBP12^F36V^-eGFP, via HaloFK7, induces the targeted hyperacetylation of eGFP. To validate that the acetyltransferase domain of HaloTag7-p300^1048–1660^ is responsible for the increase in eGFP’s acetylation status, a catalytically inactive HaloTag7-p300(I1395R)^1048–1660^ fusion protein was created and subjected to the same conditions as the catalytically active protein. Similarly to HaloTag7-p300^1048–1660^, we observe the presence of HaloTag7-p300(I1395R)^1048–1660^ within the GFP pulldown only in the presence of HaloFK7 (Fig. 2F). However, we do not observe an increase in the acetylation status of eGFP upon ternary complex formation with the catalytically dead variant (Fig. 2E). Additionally, we do not observe auto-acetylation of p300, a key indicator of its activity, thus verifying the loss of activity in our HaloTag7-p300(I1395R)^1048–1660^ mutant (Fig. S3). This data establishes that the ternary complex with active p300 is responsible for alterations in eGFP’s acetylation status. Concurrently with immunoblotting experiments, we performed mass spectrometry analysis to identify eGFP specific lysine residues and their relative acetylation abundance upon ternary complex formation (Supplemental Data 1). We observed an increase in acetylation at K156 and K158, located on an exposed loop between beta strands 7 and 8, in the presence of HaloFK7 ^40, 41^.

**Figure 2:**
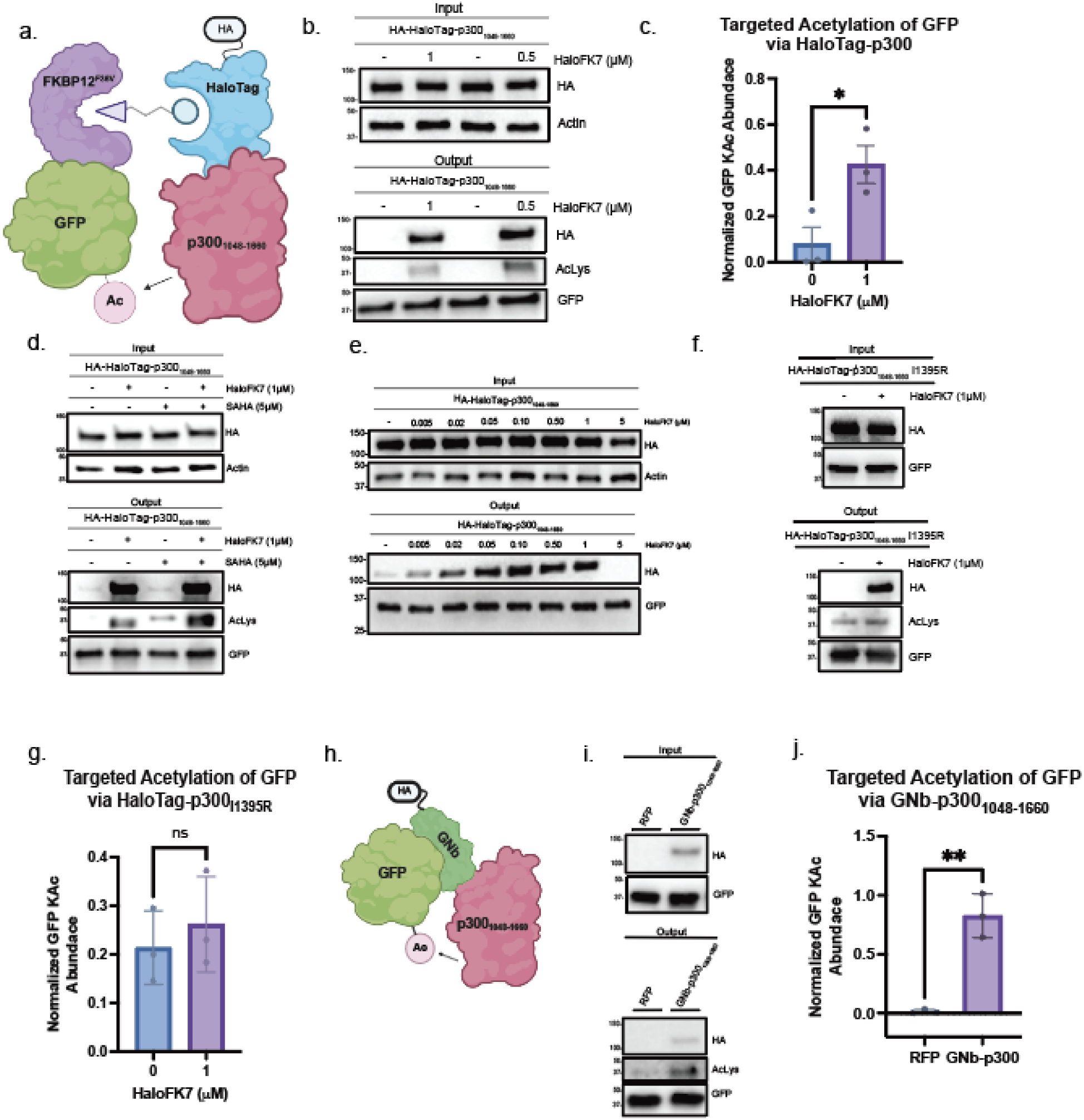
Proof of Concept - Targeted acetylation of eGFP by p300. A) Scheme depicting ternary complex formation of fusion proteins FKBP12^F36V^-eGFP and HaloTag7-p300_1048-1660_ by heterobifunctional molecule, HaloTagFK7. B) GFP Trap pull down western blot of lysine acetylation for FKBP12^F36V^-eGFP at varying concentrations of HaloFK7. HaloTag7-p300_1048-1660_ was transiently transfected into FKBP12^F36V^-eGFP cells and treated with either DMSO or HaloFK7 (1μM and 0.5μM) for 18 hrs, lysed, and FKBP12^F36V^-eGFP was enriched and blotted with indicated antibodies. C) Quantification of lysine acetylation (Kac) of eGFP relative to FKBP12^F36V^-eGFP by HaloTag7-p300_1048-1660_ as the mean ± sem for n=3 biological replicates. D) GFP Trap pull down western blot of lysine acetylation for FKBP12^F36V^-eGFP in the presence of the deacetylase inhibitor SAHA. SAHA positive cells were treated with 5μM SAHA 2 hrs prior to harvesting. E) Western blot of compound-dependent ternary complex formation of FKBP12^F36V^-eGFP and HaloTag7-p300_1048-1660_. F) GFP Trap pull down western blot of lysine acetylation for FKBP12^F36V^-eGFP by catalytically inactive HaloTag7-p300_1048-1660_I1395R. G) Quantification of lysine acetylation (Kac) of eGFP relative to FKBP12^F36V^-eGFP by HaloTag7-p300_1048-1660_I1395R as the mean ± sem for n=3 biological replicates. H) Scheme depicting binary complex formation of nanobody fusion protein GNb-p300_1048-1660_ to eGFP. I) GFP Trap pull down western blot of lysine acetylation for eGFP by GNb-p300_1048-1660_. J) Quantification of lysine acetylation (Kac) of eGFP by GNb-p300_1048-1660_ as the mean ± sem for n=3 biological replicates.

To demonstrate the compound independent mode, we replaced the HaloTag with a GFP Nanobody (GNb) using our modular cloning strategy, thus creating GNb-p300^1048–1660^, and transfected it into FKBP12^F36V^-GFP cells. Due to its high affinity and selectivity to eGFP, GNb maintains the complex formation previously observed by HaloTag7-p300^1048–1660^ to GFP for targeted acetylation (Fig. 2H). We only observed the formation of a binary complex and an increase in the acetylation of eGFP in the presence of GNb-p300^1048–1660^ (Fig. 2I, 2J). The validation of nanobodies using our platform offers significant advantages, as they exhibit high stability, strong target affinity, minimal cytotoxicity, and ability to be used without tagging the protein of interest thus broadening the range of potential applications for this approach. These results indicate that we have established a method of altering the acetylation status of the neosubstrate, eGFP, utilizing either small molecule or nanobody induced proximity.

### Expanding the Suite of Acetyltransferases for Targeted Histone Acetylation

One advantage of our modular strategy is the ability to rapidly test lysine acetylation specificity through various acetyltransferases for different proteins of interest. Histone 3 (H3) is a core component of the nucleosome, a multi-protein complex wrapped in DNA, allowing the formation of chromatin ^42, 43^. Site-specific acetylation of the H3-tail and its corresponding acetyltransferases have been widely characterized due to their involvement in gene regulation ^1, 31, 43, 44^. To evaluate our platform’s ability to incite targeted acetylation beyond cytosolic proteins, we used our HA-HaloTag7-p300^1048–1660^ expression vector and transfected it into HEK293T cells stably transduced with a FKBP12^F36V^-H3 fusion construct (Fig. 3A, Fig S1). We utilized site-specific H3 acetylation antibodies to evaluate the changes in H3-tail acetylation induced by HaloFK7. For HaloTag7-p300^1048–1660^, we observed an increase in H3K9Ac and H3K27Ac, known lysine targets of p300, only in the presence of HaloFK7 (Fig. 3B). This result validates our platform’s ability to acetylate nuclear proteins in a heterobifunctional dependent fashion.

**Figure 3:**
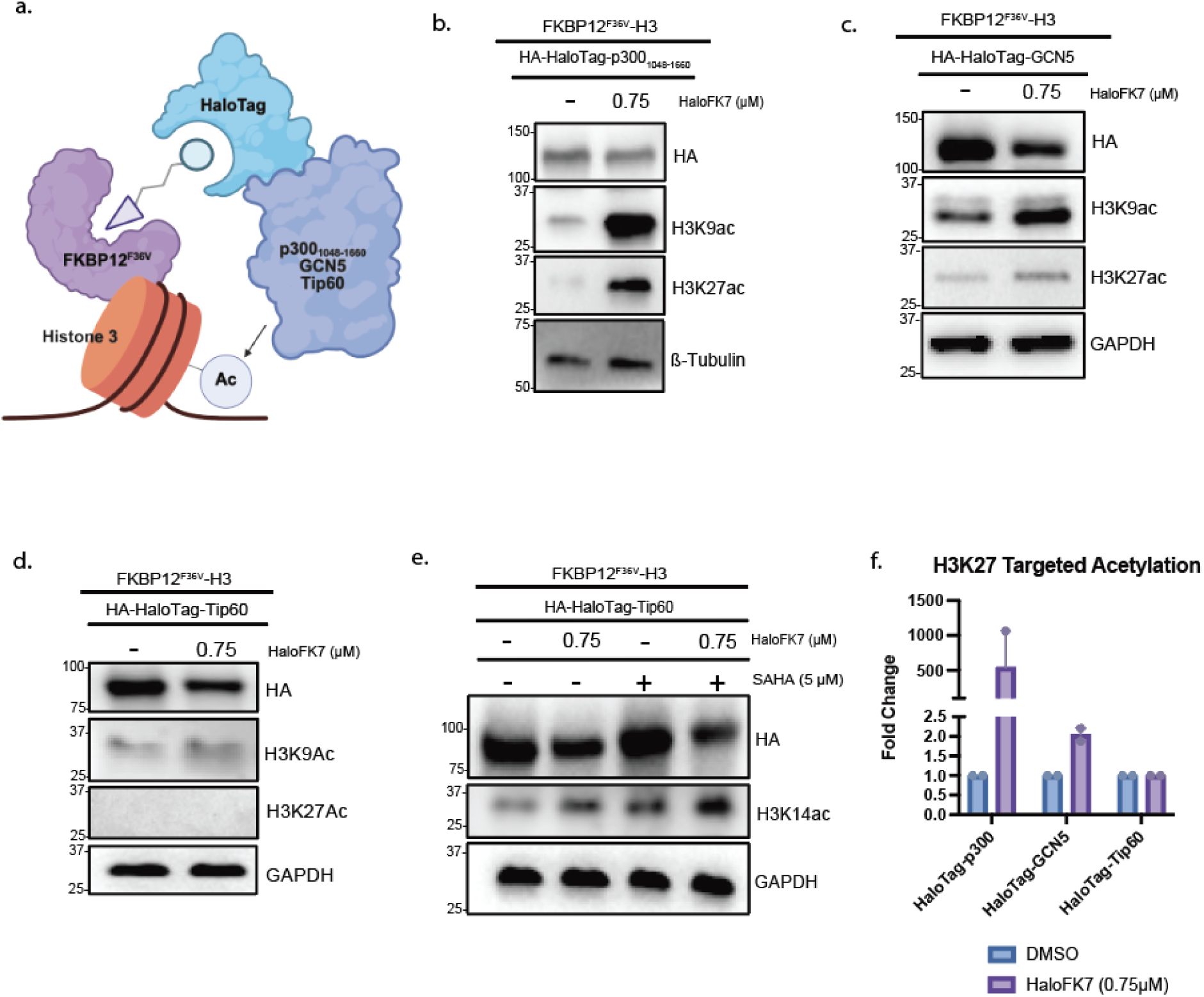
Expanding the Suite of Acetyltransferases for Targeted Histone Acetylation. A) Scheme depicting ternary complex formation of fusion proteins FKBP12^F36V^-H3 and either HaloTag7-p300_1048-1660_, HaloTag7-GCN5, or HaloTag7-Tip60 by heterobifunctional molecule, HaloTagFK7. B) GFP Trap pull down western blot of H3K9Ac, H3K27Ac for FKBP12^F36V^-H3 in the presence of HaloFK7 by HaloTag7- p300_1048-1660._ HaloTag7-p300_1048-1660_ was transiently transfected into FKBP12^F36V^-H3 cells and treated with either DMSO or HaloFK7 (0.75 μM) for 5 hrs, lysed, and blotted with noted antibodies. C) GFP Trap pull down western blot of H3K9Ac, H3K27Ac for FKBP12^F36V^-H3 in the presence of HaloFK7 by HaloTag7-GCN5_._ D) GFP Trap pull down western blot of H3K9Ac, H3K27Ac for FKBP12^F36V^-H3 in the presence of HaloFK7 by HaloTag7-Tip60._._ E) GFP Trap pull down western blot of H3K14ac for FKBP12^F36V^-H3 in the presence of the deacetylase inhibitor SAHA by HaloTag7-Tip60. SAHA positive cells were treated with 5μM SAHA 1hr prior to harvesting. F) Fold change quantification of H3K27Ac of FKBP12^F36V^-H3 by HaloTag7-p300_1048-1660_, HaloTag7-GCN5, HaloTag7-Tip60 as the mean ± sem for n=2 biological replicates.

Using this substrate, we further verified the platform’s modularity by rapidly varying effector domains. We constructed HA-HaloTag7-GCN5 and HA-HaloTag7-Tip60 expression vectors, other acetyltransferases known to interact with the H3-tail. Expression of the GCN5 construct also resulted in an increase in H3K9Ac and H3K27Ac, known lysine targets of GCN5, only in the presence of HaloFK7 (Fig. 3C). Crucially by using GCN5, we can chemically induce the acetylation of H3 at known sites without the broad proteome wide changes in acetylation associated with p300 overexpression (Fig. S2). In contrast, we don’t observe an increase in H3K27Ac for Tip60, as H3K27 is not a Tip60 substrate (Fig. 3D). However, we do observe an increase in H3K9Ac and H3K14Ac, which are known Tip60’s H3 substrates (Fig. 3D, 3E) ^45–48^. Additionally, we observed variations in the magnitude of acetylation for H3K27 (Fig. 3F). Combined, this data suggests targeted acetylation is dictated by the acetyltransferase’s substrate preference and catalytic turnover rather than induced proximity alone. Overall, these results show that our modular system can be readily adapted to different acetyltransferases, which can be used to tailor the histones marks deposited, validating the platform’s use for site-specific acetylation in a chromatin context.

### Targeted Acetylation of p53 by Xeno PTM Editor

Like Histone 3, the acetylation state of p53 is highly characterized as it plays a critical role in its activation, regulation, and interaction with other biomolecules and thus, p53’s key tumor-suppressing downstream effects ^12, 17, 49–51^. To assess the tool’s ability to alter p53 acetylation, we created a FKBP12^F36V^-p53 stable cell line (Fig S1). In the presence of HaloTag7-p300^1048–1660^, we observed an increase in the acetylation of FKBP12^F36V^-p53 K382 only when cells are treated with HaloFK7 (Fig. 4A, 4B). Additionally, we observed negligible p53K382Ac changes in endogenous p53 when compared to FKBP12^F36V^-p53 highlighting the selectivity of our platform to primarily target our engineered fusion proteins. Since p300 can likely acetylate other lysine residues within and outside of p53’s C-terminal domain, we employed mass spectrometry and identified multiple lysine residues within the C-terminal domain as well as DNA binding domain that displayed an increase in acetylation upon treatment with HaloFK7 (Supplemental Data 2).

**Figure 4:**
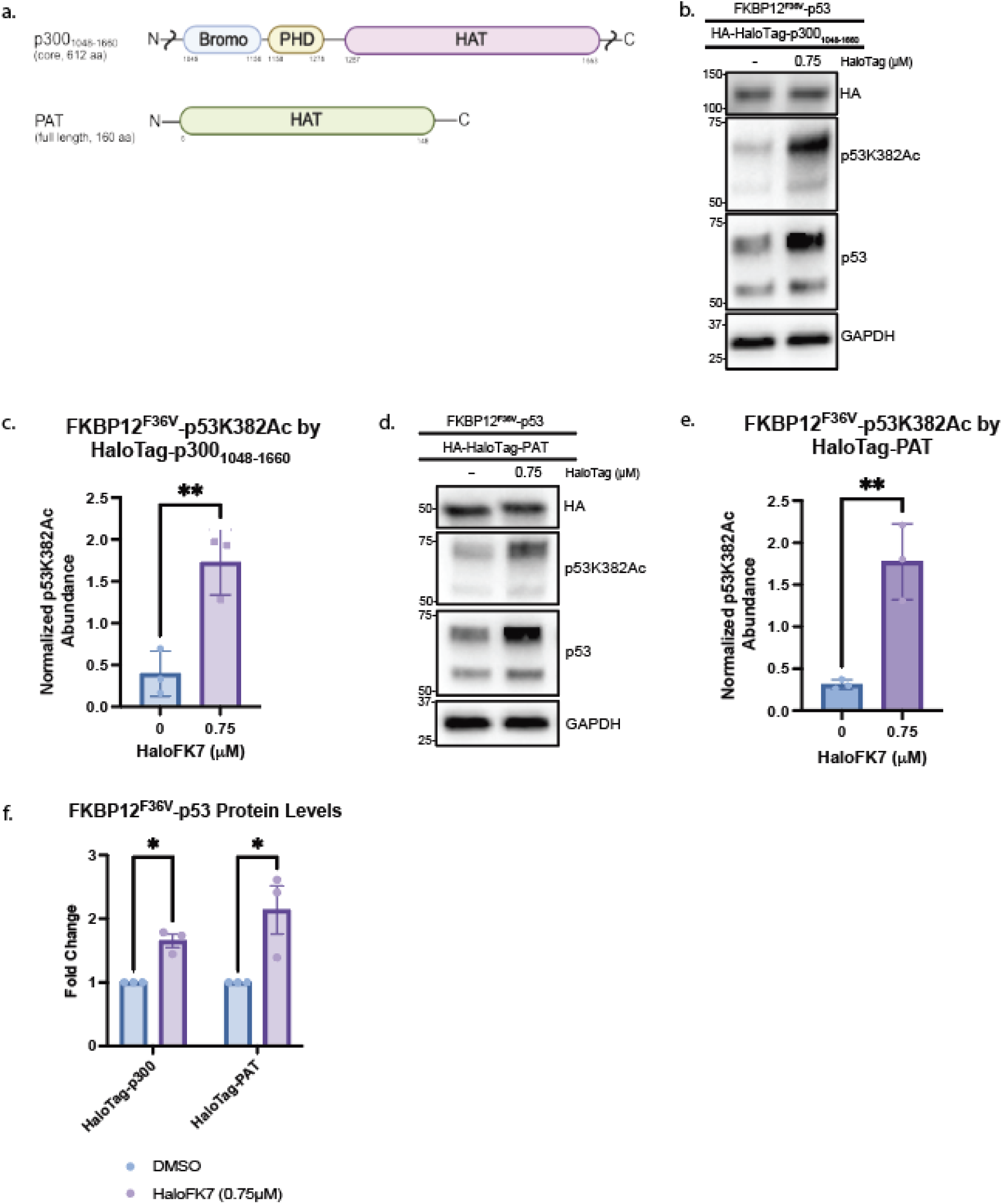
Targeted Acetylation of p53 by Xeno PTM Editor. A) Scheme depicting domain variations for HaloTag7-p300_1048-1660_ and HaloTag7-PAT B) Western blot of p53K382Ac for FKBP12^F36V^-p53 and p53 WT in the presence of HaloFK7 by HaloTag7-PAT. HaloTag7-p300_1048-1660_ was transiently transfected into FKBP12^F36V^-p53 cells and treated with either DMSO or HaloFK7 (0.75 μM) for 5 hrs, lysed, and blotted with noted antibodies. C) Quantification of p53K382Ac relative to FKBP12^F36V^-p53 by HaloTag7-p300_1048-1660_ as the mean ± sem for n=3 biological replicates. D) GFP Trap pull down western blot of p53K382Ac for FKBP12^F36V^-p53 and p53 WT in the presence of HaloFK7 by HaloTag7-PAT. E) Quantification of p53K382Ac relative to FKBP12^F36V^-p53 by HaloTag7-PAT as the mean ± sem for n=3 biological replicates. F) Quantification of change in FKBP12^F36V^-p53 protein abundance induced by HaloTag7-p300_1048-1660_ and HaloTag7-PAT in the presence of HaloFK7, as the mean ± sem for n=3 biological replicates.

As we explored the platform’s capabilities, we sought to further address the proteome-wide promiscuity of p300 and identify a way to acetylate desired lysine residues while leaving the remaining proteome unaffected*. Sulfolbus solfataricus* protein acetyltransferase (PAT) is a small (18.6 kD) archaeal acetyltransferase that shares an acetyl-CoA binding core with the much larger eukaryotic HATs (Fig. 4C) ^52^. We generated a HA-HaloTag7-PAT expression vector and transfected it into HEK293T cells stably transduced with a FKBP12^F36V^-p53 fusion construct to assess whether PAT could acetylate p53, in a similar manner to p300, but without impacting the rest of the proteome. We again observed an increase in the acetylation of FKBP12^F36V^-p53K382 in the presence of HaloFK7 (Fig. 4D, 4E). Congruently with HaloTag7-p300^1048–1660^, we observe an overall increase in the abundance of FKBP12^F36V^-p53 in the presence of the HaloFK7 (Fig. 4F), likely due to inhibition of the binding of the E3 ligase MDM2 ^49, 50, 53, 54^. This data suggests that acetylation induced by PAT can stabilize FKBP12^F36V^-p53 in a mechanistically similar fashion to p300 but with a significantly decreased amount of promiscuous acetylation with PAT expression when compared to p300 (Fig. S2). Additionally, via mass spectrometry we detected a similar profile of lysine acetylation on FKBP12^F36V^-p53 with HA-HaloTag7-PAT as to HA-HaloTag7-p300 upon treatment with HaloFK7 (Supplemental Data 3). HaloTag7-PAT enables us to perform targeted acetylation of p53 with minimal off-target effects, introducing the concept and value of a xeno-PTM editor for targeted protein acetylation. Collectively, these results demonstrate that our modular platform dependably recapitulates native acetyltransferase substrate specificity, enabling precise and programmable site-specific acetylation of various proteins throughout the cell and therefore guides the development of heterobifunctional compounds.

## Conclusion

Here, we’ve established and validated a modular PTM editing platform that can perform targeted acetylation on various proteins of interest, further expanding the biochemical tools currently available for studying PTMs. We demonstrated we can alter the acetylation status of eGFP, histone 3, and p53, using either compound-dependent or nanobody based recruitment of acetyltransferases. Starting with p300, due to its promiscuity and known use with neo-substrates ^11, 37–39, 55, 56^, we validated our platform’s intracellular PTM editing capabilities against varying substrates. For each substrate we tested, p300 was successfully localized to the substrate and able to hyperacetylate available lysine residues. While we were able to build the foundation and perform targeted acetylation at a substrate-specific level, we were concerned by the proteome-wide acetylation increases observed by p300 overexpression.

We thus expanded and validated the platform’s ability to rapidly assemble fusion proteins varying the acetyltransferase effector domains. We broadened the accessibility to other subfamilies of acetyltransferases specifically the GNAT and MYST families. GNAT and MYST acetyltransferase overlap functionally through their ability to form the covalent linkage between acetyl-coA and lysine but structurally they differ with GNATs possessing a bromodomain essential for acetylated lysine recognition in contrast to the MYST family zinc finger domain^2, 44, 57^. Both families possess higher substrate specificity when compared to p300 while still retaining enzymatic activity. Using HaloFK7, we were able to induce proximity of GCN5 (GNAT) and Tip60 (MYST) to FKBP12^F36V^-H3 and FKBP12^F36V^-p53, thereby altering their acetylation status. Finally, we sought to further diversify the platform by validating its ability to use non-mammalian acetyltransferase for future biological exploration ^52^. The archaeal acetyltransferase, PAT, was successfully recruited to FKBP12^F36V^-p53 via HaloFK7, and able to mimic p300-induced FKBP12^F36V^-p53 hyperacetylation while limiting proteome-wide changes. Through our findings, we discovered that site-specific acetylation is dictated by the acetyltransferase recruited to the protein of interest rather than the platform’s design, thus addressing the shortcomings of using p300 alone.

We recognize that further optimization may be required, including but not limited to linker length and composition within the fusion proteins. However, the user-friendly modular design provided by this approach allows for this tunability to occur with precision and ease. Our technology can rapidly provide proof of concept and information about the ideal effector for either a heterobifunctional molecule or a nanobody-fusion design for further studies, addressing a gap with current biological tools to study PTMs.

In conclusion, we present a modular, user-friendly toolbox for editing PTMs, exemplified by acetylation. Using this approach, we demonstrate nanobody and heterobifunctional compound induced acetylation with a variety of acetyltransferases, against a suite of substrates, demonstrating that the choice of transferase can dictate the acetyl marks deposited. Additionally, we introduce the concept of xeno-PTM editor recruitment to reduce off-target PTM editing in mammalian cells as well as effective nanobody targeted acetylation, further expanding, and increasing accessibility to, the biochemical toolbox of studying PTMs.

## Acknowledgements

We thank members of the Burslem lab for comments and intellectual discussion over the course of this study. We would also like to thank Drs Ronen Marmorstein, Erica Korb, Mark Sellmyer and Vikram Paralkar for support and guidance during this project. This work was supported by the National Institute of General Medical Sciences (R35GM142505) to G.M.B., the National Institute of Diabetes and Digestive and Kidney Diseases (T32DK007780) and by the Howard Hughes Medical Institute (HHMI) Gilliam Fellowship for Advanced Study to B.H.P.

## Supporting Information

The supporting information contains Supplemental Figure (S1-3) and Tables (S1), mass spectrometry data, as well as a detailed protocol for 3-component Gibson assembly and other materials and methods.

## Notes

### Competing Interest Statement

The authors have declared no competing interest.

