## Supplemental Information for "A Modular Platform for Effector Discovery in Induced-Proximity Lysine Acetylation"

**Supplemental Figures**

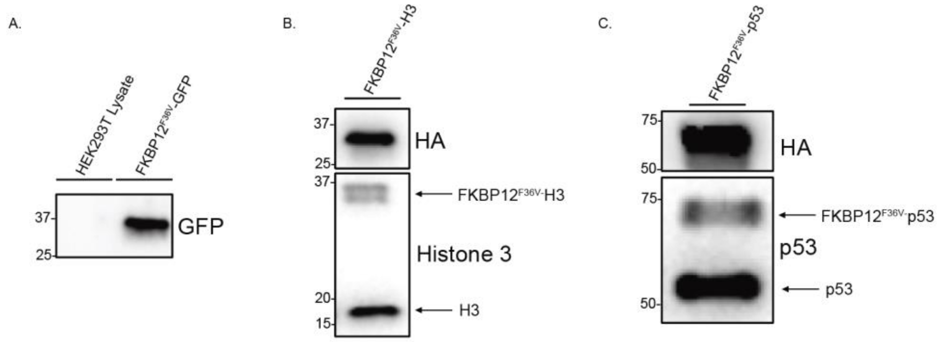

**Figure S1. Validation of model cell lines (FKBPV-eGFP (A), FKBPV-Histone3 (B), FKBPV-p53 (C))**

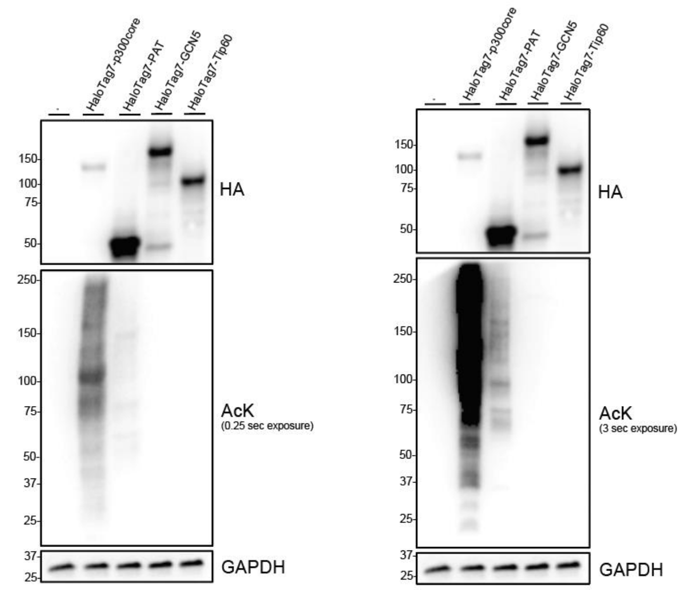

**Figure S2 – Proteome wide alterations in acetylation with effector overexpression. Pan acetyl lysine blots with short exposure (left) and long exposure (right) of cells overexpressing HaloTag-p300, HaloTag-GCN5, HaloTag-PAT, and HaloTag-Tip60.**

**Table S1 Library of Fusion Protein Vectors created via Modular PTM editing platform**

| | |
|---|---|
| 1 | Flag-HaloTag-p300 (pcDNA5) |
| 2 | HA-HaloTag-p300 (pcDNA5) |
| 3 | HaloTag-p300-His (pcDNA5) |
| 4 | HA-HaloTag-p300-I1395R (pcDNA5) |
| 5 | HA-GNb-p300 (pcDNA5) |
| 6 | HA-HaloTag-GCN5 (pcDNA5) |
| 7 | HA-HaloTag-Tip60 (pcDNA5) |
| 8 | HA-FKBP$^{F36V}$-p53 (Lenti) |
| 9 | HA-FKBP$^{F36V}$-H3 (Lenti) |
| 10 | HA-HaloTag-HDAC6 (pcDNA5) |
| 11 | HA-HaloTag-Sirt1 (pcDNA5) |
| 12 | HA-FKBP$^{F36V}$-GFP-X3 (Lenti) |

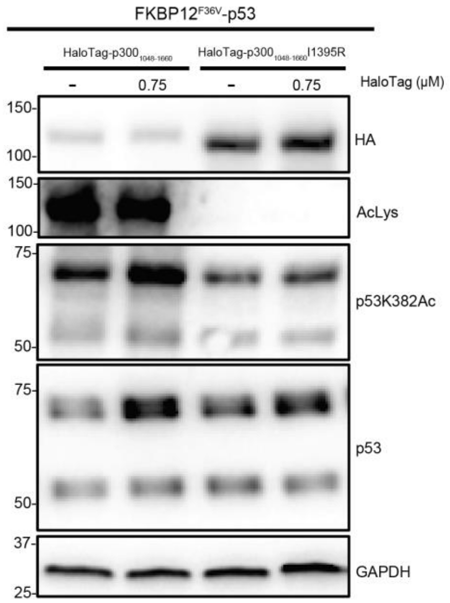

**Figure S3 - Targeted acetylation of FKBP$^{F36V}$-p53 is only observed when catalytically active HaloTag-p300$^{1048\text{-}1660}$ is recruited, not inactive (I1395R) HaloTag-p300$^{1048\text{-}1660}$**

**Detailed Protocol for Optimized Fusion Protein Plasmid Generation**

Below is a step-by-step guide for the construction of HA-HaloTag-p300$^{1048\text{-}1660}$ and the same method was used to construct all fusion proteins within our modular PTM editing platform.

Step 1: Development of Modular Gibson Constructs (Targeting, Effector, and Backbone Domains)

1. Design Gibson construct primers
    a. Modular overhangs will need to be installed/amplified onto each component (targeting, effector domain, effector domain, and backbone)
    b. Calculation of Tm should only involve section of the primer that is on parent plasmid, not the modular overhangs.

c. The overhangs listed below create a modular platform where each fusion protein contains an N-terminal HA epitope tag and a $(GlySer_4)_2$ linker.

Commented [GB1]: Lets provide a detailed step-by-step guide in the SI with photos etc.

Targeting Domain Overhangs (N Term HA Tag)

i. N-term: 5'-ATGTATCCATACGACGTTCCTGATTATGCG-3'

ii. C-term: 5'- TCCGGAACCTCCGCCACCACTACCGCCTCCACCAGA-3'

Effector Domain Overhangs

iii. N-term: 5'- TCTGGTGGAGGCGGTAGTGGTGGCGGAGGTTCCGGA-3'

iv. C-term: 5'- AGATGGCTAGAGCGTACGTTACCGAGCCTG-3'

Expression Backbone

v. N-Term: 5'- CAGGCTCGGTAACGTACGCTCTAGCCATCT-3'

vi. C-Term: 5'- CGCATAATCAGGAACGTCGTATGGATACAT-3'

d. Epitope tags, their terminal positions, and linker composition can all be customized for the need of the experiment during primer design.

2. PCR Amplification of Gibson Constructs

a. Create 10 µM stock of primers

b. Create 1 ng stock of plasmids

c. Create the following master mix on ice:

| | |
|---|---|
| Nuclease Free (NF) Water | 28.5 µL |
| Plasmid (1 ng) | 3 µL |
| FWD Primer | 3 µL |
| REV Primer | 3 µL |
| Q5 Master Mix | 37.5 µL |

d. Aliquot master mix into three 25 µL PCR tubes

e. Run the following PCR cycle for each plasmid:

pcDNA5/FRT (5,070 bases) Tm=65 ºC

| Temperature | Time | Cycle |
|---|---|---|
| 98º | 3 min | |
| 98º | 10 sec | 35 X |
| 60-70º | 30 sec | |
| 72º | 2 min 45 sec | |
| 72º | 2 min | |
| 4º | ∞ | |

p300 Core (1,851 bases) Tm= 63 ºC

| Temperature | Time | Cycle |
|---|---|---|
| 98º | 3 min | |
| 98º | 10 sec | 35 X |
| 58-68º | 30 sec | |
| 72º | 1 min | |
| 72º | 2 min | |

| | |
|---|---|
| 4º | ∞ |

HaloTag (891 bases) Tm=67 ºC

| Temperature | Time | Cycle |
|---|---|---|
| 98º | 3 min | |
| 98º | 10 sec | 35 X |
| 62-72º | 30 sec | |
| 72º | 30 sec | |
| 72º | 2 min | |
| 4º | ∞ | |

f. Pour DNA gel

g. Add 5 μL of DNA loading dye to the each 25 μL PCR reaction after the cycle is complete

h. Load 25 μL of sample into each well and 5 μL of DNA ladder

i. Run gel in 1X Tris-acetate-EDTA (TAE, 40mM Tris-acetate, 1mM EDTA, pH 8.0) buffer at 130V

j. Excise bands

k. Perform gel extraction (QIAquick Gell Extraction Kit Cat#: 28706) to retrieve Gibson constructs

l. Measure DNA concentration of each construct

Step 2: Multicomponent Gibson Assembly

1. Complete the calculations for constructs using NEB Gibson Assembly Calculator
    a. 3:1 ratio was using for mammalian vectors. However, 2:1 ration was used for lentiviral vector due to the larger overall size.

| | 3:1 |
|---|---|
| p300 Core (1,913 bases) | 100.1 ng |
| HaloTag (891 bases) | 50.05 ng |
| pcDNA5/FRT (5,070 bases) | 83.42 ng (27 fmol) |
| Nuclease Free (NF) Water | 10-total constructs volume μL |
| 2X Master Mix | |

2. Gibson Assembly
    a. Add respected amounts to PCR tube
    b. 60 min at 50 ºC then 4 ºC infinite
3. Transformation
    a. Add 3 μL of Gibson reaction to DH5α competent cells or NEB-5α competent cells
    b. Mix contents

c. Incubate on ice for 30 min
d. Heat shock at 42 ºC for 45 sec
e. Incubate on ice for 2 min
f. Add 600 μL of SOC media
g. Shake at 37 ºC for 1 hour
h. Plate on Amp plate for pcDNA5/FRT/TO backbone
i. Place plate in 37 ºC incubator overnight
j. Select colonies for grow up and place in 5 mL of LB Broth with 5 μL of Amp antibiotic & placed in 37 ºC shaker overnight
k. Perform plasmid extraction
l. Take plasmid concentration and sequence for verification

*PCR Amplification of Gibson Assembly Constructs*

NEB Q5® High Fidelity 2X Master Mix or Phusion™ High Fidelity PCR Master Mix with HF Buffer was used to PCR amplify targeting domains, effector domains and backbones with modular overhangs. Primers were created to include full-length overhangs as well as amplified protein.

Targeting Domain Overhangs

N-term: 5'-ATGTATCCATACGACGTTCCTGATTATGCG-3'

C-term: 5'- TCCGGAACCTCCGCCACCACTACCGCCTCCACCAGA-3'

Effector Domain Overhangs

N-term: 5'- TCTGGTGGAGGCGGTAGTGGTGGCGGAGGTTCCGGA-3'

C-term: 5'- AGATGGCTAGAGCGTACGTTACCGAGCCTG-3'

Expression Backbone

N-Term: 5'- CAGGCTCGGTAACGTACGCTCTAGCCATCT-3'

C-Term: 5’- CGCATAATCAGGAACGTCGTATGGATACAT-3'

#### Materials and Methods

##### Gibson Assembly

A three-component Gibson assembly was performed by the addition of constructs (targeted domain, effector domain or protein of interest, and expression backbone) at ratios depending upon their size and NEBuilder® HiFi DNA Assembly Master Mix. All components were heated to 50 °C for one hour. Gibson assembly product was transformed in NEB 5α Competent E.coli (High efficiency) competent cells. Plasmids were extracted via QIAprep Spin Miniprep Kit and verified by whole plasmid sequencing via Plasmidasaurus.

##### Generation of Stable Cell Lines

Stable cell lines were created for the following target proteins: FKBP12$^{F36V}$-eGFP, FKBP12$^{F36V}$-p53, FKBP12$^{F36V}$-H3. Lentiviral vectors containing the target protein fused to the FKBP12$^{F36V}$ protein were generated via a modular cloning strategy. The virus was generated through the transduction of viral packaging agents, pmD2.G and PAX2, and a lentiviral vector into human embryonic kidney (HEK) 293 cells. Virus was harvested and concentrated using Abcam Virus Precipitation Kit (ab102538). Concentrated virus was added to human embryonic kidney (HEK) 293 cells or Flp-In$^{TM}$ T-REx 293 cells, followed by Puromycin or Puromycin/Zeocin selection to generate a stable expressing cell line of the targeted fusion protein.

##### Cell Culture

Human embryonic kidney (HEK) 293 cells were cultured in Dulbecco’s Modified Eagle Medium (DMEM) containing 5% Penicillin/Streptomycin and 10% Fetal Bovine Serum (FBS). Flp-In$^{TM}$ T-REx 293 cells were cultured in Dulbecco’s Modified Eagle Medium (DMEM) containing 5% Penicillin/Streptomycin, 10% Fetal Bovine Serum (FBS).

##### Western Blotting

Cells were lysed in lysis buffer (20 mM Tris-HCl (pH 7.5), 150 mM NaCl, 1 mM Na2EDTA, 1 mM EGTA, 1% NP-40, 1% sodium deoxycholate, 2.5 mM sodium pyrophosphate, 1 mM beta-glycerophosphate, 1 mM $Na_3VO_4$, 1 µg/ml leupeptin) supplemented with protease and deacetylase inhibitors (Thermo Scientific Pierce™ Protease Inhibitor Mini Tablets, EDTA-Free; Cat#: A32955 and Med Chem Express Deacetylase Inhibitor Cocktail; Cat #: HY-K00300). The protein concentrations of whole-cell lysates were determined with the Pierce™ Rapid Gold BCA Protein Assay Kit. 20 µg of whole-cell lysates were resolved by SDS-PAGE and transferred to nitrocellulose membranes (Bio-Rad). Membranes were blocked with 5% non-fat dry milk in TBST (Tris-buffered saline with 0.05% TWEEN-20, pH 8.0) and probed with indicated antibodies. Immunoreactive proteins were visualized using enhanced chemiluminescence detection substrate (Amersham ECL Prime Western Blotting Detection Reagent RPN2232) using a Molecular Imager ChemiDoc XRS+ System (BioRad). The following primary antibodies were used for this manuscript:

- Acetylated Lysine (Cell Signaling Technologies Cat#: 9441S)
- GFP (Cell Signaling Technologies Cat#: 2956S)
- HA Tag (Cell Signaling Technologies Cat#: 3724S)
- Histone 3 (Cell Signaling Technologies Cat#: 4499S)
- Acetyl-histone 3 (H3K9) (Cell Signaling Technologies Cat#:9649S)
- Acetyl-histone 3 (H3K14) (Millipore Cat#: 07-373)
- Acetyl-histone 3 (H3K27) (Cell Signaling Technologies Cat#: 8173S)
- p53 (Cell Signaling Technologies Cat#: 2524S)
- Acetyl-p53 (K382) (Cell Signaling Technologies Cat#: 2524S)
- GAPDH (Cell Signaling Technologies Cat#: 5174S)
- Actin (Cell Signaling Technologies Cat#: 8457S)

**Targeted Acetylation via Heterobifunctional Molecule**

Cells stably expressing target protein were seeded (~0.3e6 cells per well) in 6-well plates. Mammalian expressing vector, containing effector fusion protein, was transfected into a stable expressing target protein cell line using PolyJet™ In Vitro DNA Transfection Reagent. Heterobifunctional compound, HaloFK7, or DMSO was added to cells and incubated for ~18 hours. Cells were harvested and lysed with 1X RIPA buffer (Cell Signaling Technology Cat.#: 9806 - 20 mM Tris-HCl (pH 7.5), 150 mM NaCl, 1 mM $Na_2EDTA$, 1 mM EGTA, 1% NP-40, 1% sodium deoxycholate, 2.5 mM sodium pyrophosphate, 1 mM beta-glycerophosphate, 1 mM $Na_3VO_4$, 1 µg/ml leupeptin) containing protease (Thermo Scientific Pierce™ Protease Inhibitor Mini Tablets, EDTA-Free; Cat#: A32955) and deacetylase (Med Chem Express Deacetylase Inhibitor Cocktail; Cat #: HY-K00300) inhibitors. After clearing the lysates, protein concentration was determined using the Pierce™ Rapid Gold BCA assay. Lysate concentration was standardized. For GFP: pull-down was performed on standardized lysates via ChromoTek GFP-Trap® Agarose, followed by immunoblot using pan-acetyl lysine to assess the acetylation status of GFP. Acetyl-lysine specific antibodies were used to assess the acetylation status of p53 and H3 whole cell lysates via immunoblot.

**Targeted Acetylation via Nanobody**

Cells stably expressing target protein were seeded (~0.3e6 cells per well) in 6-well plates. Mammalian expressing vector, containing effector fusion protein, was transfected into a stable expressing target protein cell line using PolyJet™ In Vitro DNA Transfection Reagent and incubated for ~18 hours. Cells were harvested and lysed with 1X RIPA buffer (Cell Signaling Technology Cat.#: 9806 - 20 mM Tris-HCl (pH 7.5), 150 mM NaCl, 1 mM $Na_2EDTA$, 1 mM EGTA, 1% NP-40, 1% sodium deoxycholate, 2.5 mM sodium pyrophosphate, 1 mM beta-glycerophosphate, 1 mM $Na_3VO_4$, 1 µg/ml leupeptin) containing protease (Thermo Scientific Pierce™ Protease Inhibitor Mini Tablets, EDTA-Free; Cat#: A32955) and deacetylase (Med Chem Express Deacetylase Inhibitor Cocktail; Cat #: HY-K00300) inhibitors. After clearing the lysates, protein concentration was determined using the Pierce™ Rapid Gold BCA assay. Lysate concentration was standardized. For GFP: pull-down was performed on standardized lysates via ChromoTek GFP-Trap® Agarose, followed by immunoblot using pan-acetyl lysine to assess the acetylation status of GFP. Acetyl-lysine specific antibodies were used to assess the acetylation status of p53 and H3 whole cell lysates via immunoblot.

**Mass Spectrometry Sample Preparation**

Bead-bound protein samples were reconstituted in 300 µL of 50 mM triethylammonium bicarbonate (TEABC). Proteins were reduced by adding freshly prepared 100 mM dithiothreitol (DTT) to a final concentration of 10 mM, followed by incubation at 60 °C for 20 min. Subsequently, freshly prepared iodoacetamide (IA) was added to a final concentration of 20 mM, and samples were incubated at room temperature in the dark for 10 min to allow alkylation of cysteine residues.

Proteins were digested by adding MS grade trypsin/Lys-C mix (Promega, catalog no. V5072)) at an enzyme to protein ratio of 1:100 (w/w), followed by incubation at 37 °C overnight with shaking. The digestion was quenched the following day by the addition of formic acid (FA) to a final concentration of 0.1% (v/v).

**Peptide Desalting Using $C_{18}$ Stage Tips**

Digested peptides were desalted using $C_{18}$ stage tips packed with Empore™ $C_{18}$ extraction disks (cat. no. 66883-U) following a standard protocol. Stage tips were conditioned with methanol and 50% (v/v) acetonitrile containing 0.1% (v/v) formic acid, equilibrated with 0.1% (v/v) formic acid, and loaded with peptides reconstituted in 3% (v/v) acetonitrile and 0.1% (v/v) formic acid. After washing with 0.1% (v/v) formic acid, peptides were eluted using 50% (v/v) acetonitrile containing 0.1% (v/v) formic acid, vacuum-dried, and stored at -20 °C until LC-MS/MS analysis.

**LC-MS/MS Analysis**

LC-MS/MS analysis was performed on a Q Exactive Hybrid Quadrupole-Orbitrap mass spectrometer coupled to a Vanquish Neo UHPLC system. Dried peptides were reconstituted in 0.1% (v/v) formic acid and separated on an analytical $C_{18}$ column (75 µm × 50 cm, 2 µm particle size, 100 Å pore size) at a flow rate of 350 nL $min^{-1}$. Peptides were eluted using a linear gradient of 4-40% solvent B (0.1% FA in 80%

ACN) over 50 min, with a total run time of 65 min. MS1 and MS2 spectra were acquired in the Orbitrap analyzer at resolutions of 70,000 and 17,500, respectively. Full MS scans were acquired over an m/z range of 375-1750, with a maximum injection time of 100 ms and an automatic gain control (AGC) target of $3 \times 10^6$ ions. Data-dependent acquisition (DDA) was performed by selecting the top 10 most intense precursor ions using a quadrupole isolation window of 2.0 m/z, with a dynamic exclusion of 20.0 s. Fragmentation was carried out using higher-energy collisional dissociation (HCD) with a normalized collision energy (NCE) of 26%. MS/MS spectra were acquired with a maximum injection time of 50 ms and an AGC target of $1 \times 10^5$ ions.

**Data Analysis**

Raw mass spectrometry data files were processed using Proteome Discoverer (version 3.0.0.757). Spectra were searched against a human protein specific FASTA sequences using the SEQUEST search algorithm. Acetylation of lysine residues and peptide N-termini (+42.011 Da), as well as methionine oxidation, were specified as dynamic modifications, while carbamidomethylation of cysteine residues was set as a static modification. Trypsin was defined as the proteolytic enzyme, allowing up to two missed cleavages. A precursor mass tolerance of 20 ppm and a fragment mass tolerance of 0.05 Da were applied. Peptide-spectrum matches (PSMs) and peptides were filtered to a false discovery rate (FDR) of 5% using a target-decoy strategy.

**Supplemental Data 1: Targeted Acetylation eGFP via HaloTag-p300 MS Data**

61.34% Coverage of eGFP

```
1   MSKGEELFTGVVPILVELDGDVNGHKFSVSGEGEGDATYGKLTLKFICTTGKLPVPWPTL
61  VTTFSYGVQCFSRYPDHMKQHDFFKSAMPEGYVQERTIFFKDDGNYKTRAEVKFEGDTLV
121 NRIELKGIDFKEDGNILGHKLEYNYNSHNVYIMADKQKNGIKVNFKIRHNIEDGSVQLAD
181 HYQQNTPIGDGPVLLPDNHYLSTQSALSKDPNEKRDHMVLLEFVTAAGITHGMDELYK
```

Green = peptides detected

| | Abundance Ratio (Sample/Control) | Abundances (Grouped): Control | Abundances (Grouped): HaloFK7 |
|---|---|---|---|
| 2X Acetyl [K156; K158] | 1.855 | 70.1 | 129.9 |

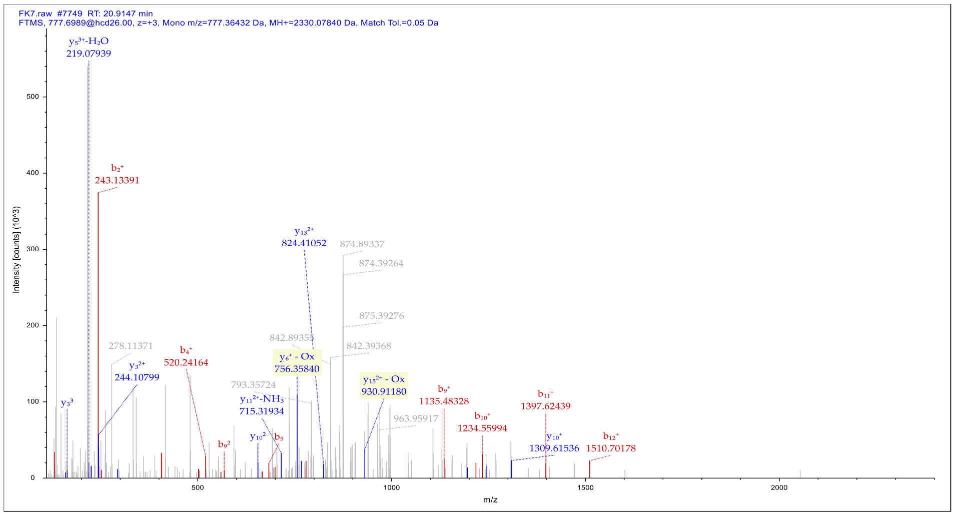

**Supplemental Data 2: Targeted Acetylation p53 via HaloTag-p300 MS Data**

94.62% Coverage of FKBP$^{F36V}$-p53

120 EEPQSDPSVEPPLSQETFSDLWKLLPENNVLSPLPSQAMDDLMLSPDDIEQWFTEDPGPD

180 EAPRMPEAAPPVAPAPAAPTPAAPAPAPSWPLSSSVPSQKTYQGSYGFRLGFLHSGTAKS

240 VTCTYSPALNKMFCQLAKTCPVQLWVDSTPPPGTRVRAMAIYKQSQHMTEVVRRCPHHER

300 CSDSDGLAPPQHLIRVEGNLRVEYLDDRNTFRHSVVVPYEPPEVGSDCTTIHYNYMCNSS

360 CMGGMNRRPILTIITLEDSSGNLLGRNSFEVRVCACPGRDRRTEEENLRKKGEPHHELPP

420 GSTKRALPNNTSSSPQPKKKPLDGEYFTLQIRGRERFEMFRELNEALELKDAQAGKEPGG

480 SRAHSSHLKSKKGQSTSRHKKLMFKTEGPDSD

Green = peptides detected (high confidence)

Yellow = peptides detected (medium confidence)

| | Abundance Ratio (Sample/Control) | Abundances : Control | Abundances : HaloFK7 |
|---|---|---|---|
| K488; K490; K491 | 18.387 | 38897.44531 | 795290.0625 |
| K499; K500 | | Not detected | 13624119 |
| K423 | | Not detected | 325298.1875 |

Modification sites: K488, K490, K491

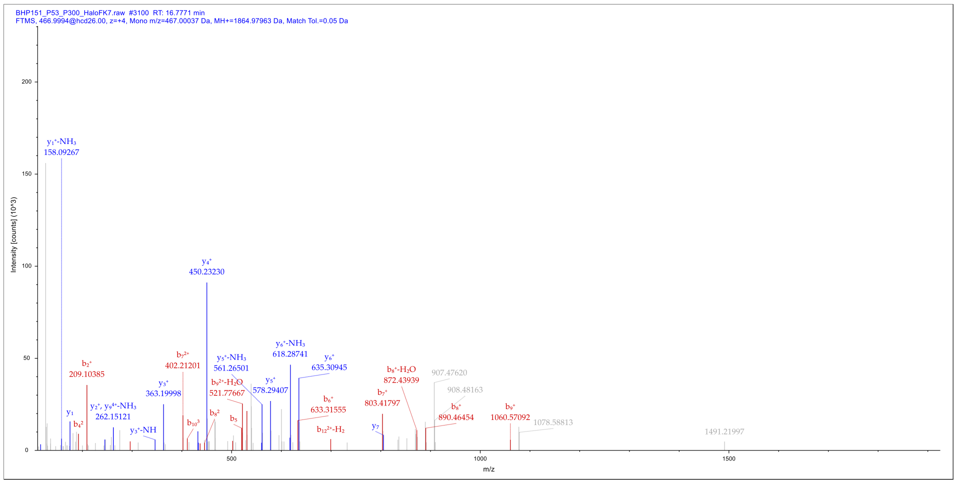

Modification sites: K499, K500

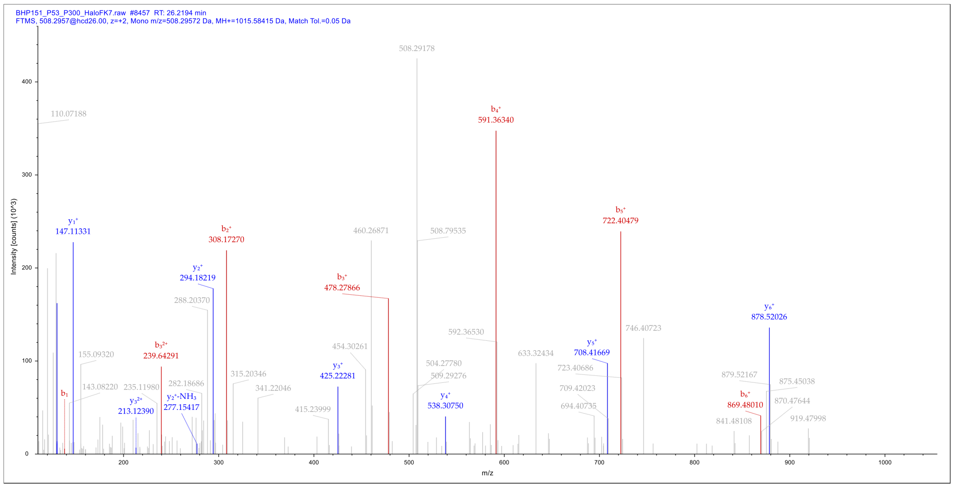

### Modification sites: K423

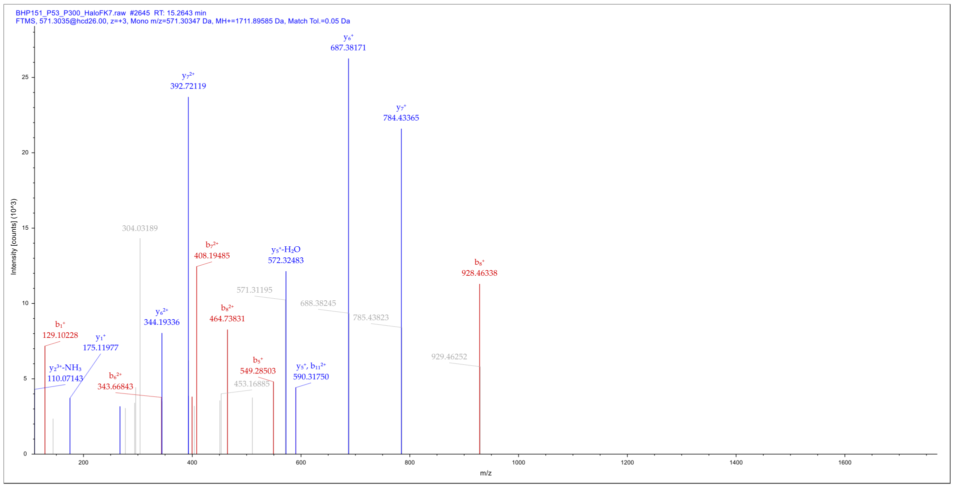

**Supplemental Data 3: Targeted Acetylation p53 via HaloTag-PAT MS Data**

93.27% Coverage of FKBP$^{F36V}$-p53

```
120  EEPQSDPSVEPPLSQETFSDLWKLLPENNVLSPLPSQAMDDLMLSPDDIEQWFTEDPGPD
180  EAPRMPEAAPPVAPAPAAPTPAAPAPAPSWPLSSSVPSQKTYQGSYGFRLGFLHSGTAKS
240  VTCTYSPALNKMFCQLAKTCPVQLWVDSTPPPGTRVRAMAIYKQSQHMTEVVRRCPHHER
300  CSDSDGLAPPQHLIRVEGNLRVEYLDDRNTFRHSVVVPYEPPEVGSDCTTIHYNYMCNSS
360  CMGGMNRRPILTIITLEDSSGNLLGRNSFEVRVCACPGRDRRTEEENLRKKGEPHHELPP
420  GSTKRALPNNTSSSPQPKKKPLDGEYFTLQIRGRERFEMFRELNEALELKDAQAGKEPGG
480  SRAHSSHLKSKKGQSTSRHKKLMFKTEGPDSD
```

Green = peptides detected (high confidence)

Yellow = peptides detected (medium confidence)

| | Abundance Ratio (Sample/Control) | Abundances : Control | Abundances : HaloFK7 |
|---|---|---|---|
| K500 | | Not detected | 1.60e5 |
| K423 | | Not detected | 2.31e5 |
| | | Not detected | 1.76e6 |

Modification Site: K500

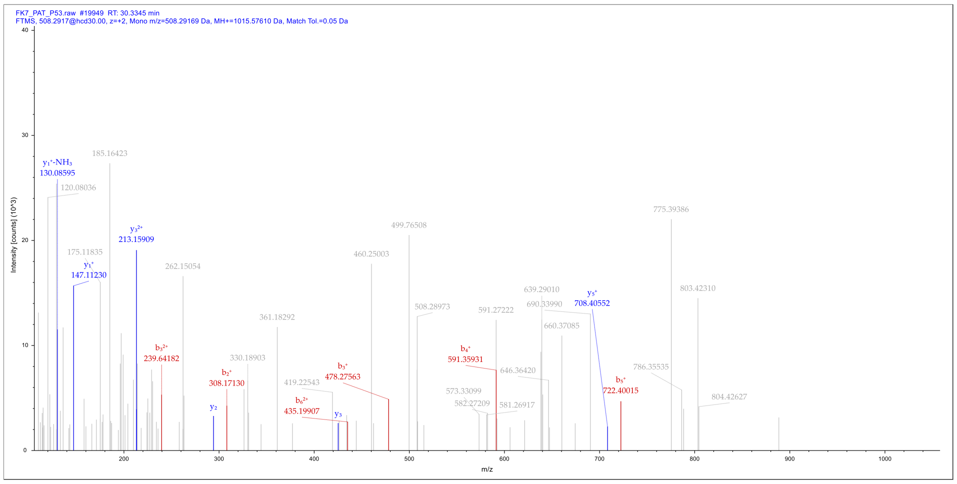

Modification Site: K423

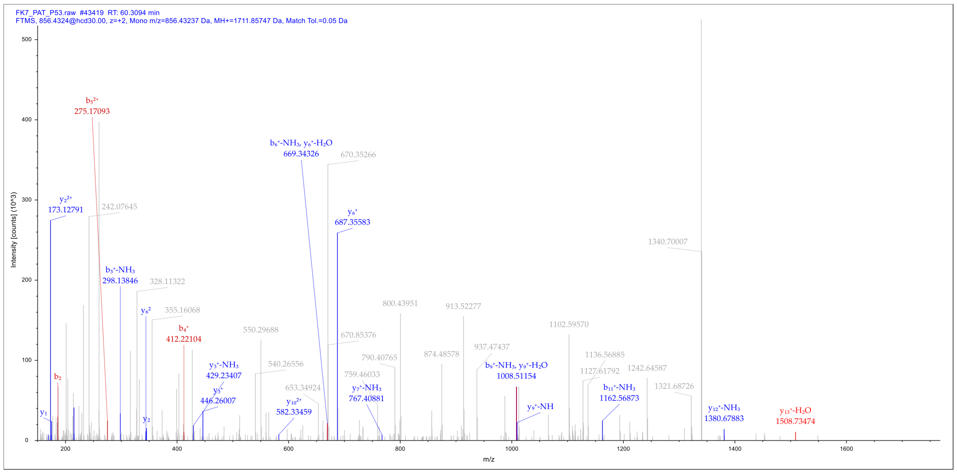

FK7_PAT_P53.raw #43419 RT: 60.3094 min
FTMS, 856.4324@hcd30.00, z=+2, Mono m/z=856.43237 Da, MH+=1711.85747 Da, Match Tol.=0.05 Da
Intensity [counts] (10^3)
m/z
$b_5^{2+}$ 275.17093
$y_2^{2+}$ 173.12791
242.07645
$b_6^+-NH_3$, $y_6^+-H_2O$ 669.34326
670.35266
$y_6^+$ 687.35583
1340.70007
$b_3^+-NH_3$ 298.13846
328.11322
$y_6^2$
355.16068
$b_4^+$ 412.22104
550.29688
800.43951
913.52277
670.85376
1102.59570
937.47437
874.48578
790.40765
1136.56885
1127.61792
1242.64587
$y_3^+-NH_3$ 429.23407
540.26556
759.46033
$b_9^+-NH_3$, $y_9^+-H_2O$ 1008.51154
$b_2$
$y_3^+$ 446.26007
653.34924
$y_7^+-NH_3$ 767.40881
1321.68726
$b_{11}^+-NH_3$ 1162.56873
$y_9^+-NH$
$y_{10}^{2+}$ 582.33459
$y_1$
$y_2$
$y_{12}^+-NH_3$ 1380.67883
$y_{13}^+-H_2O$ 1508.73474
